# Multi-vector retrieval enables residue-resolved prediction of protein partners by ColBERT-PPI

**DOI:** 10.64898/2026.09.19.752878

**Authors:** He Yang, Ruxin Lei, Youwen Zhuang, Jian Zhang

**Author notes:** Correspondence should be addressed to Y.Z. and J.Z.

## Abstract

Protein–protein interactions (PPIs) are central to biological processes, making the identification of both interacting partners and their binding sites important for understanding molecular function and guiding therapeutic discovery. However, connecting large-scale partner prediction to residue-level interaction evidence remains challenging. Here we present ColBERT-PPI, a structure-aware multi-vector framework that links protein partner retrieval to interaction-site localisation. The model retains reusable residue-level representations and compares them in a partner-dependent manner, producing raw residue compatibility maps for localisation and calibrated smooth MaxSim scores for retrieval. On PINDER, ColBERT-PPI achieved an area under the precision–recall curve of 0.482, compared with 0.151 for the strongest evaluated external comparator. Both complete and sequence-only multi-vector models outperformed the single-vector control on an independent human yeast two-hybrid screen. Residue-level comparisons highlighted experimental interfaces, captured changes in local evidence across partners and achieved a mean contact AUROC of 0.806 versus 0.526 for the single-vector control. After task-specific fine-tuning, PPI-trained representations also improved protein-RNA interaction retrieval, indicating transfer beyond the molecular context in which they were learned. By connecting candidate partners with their supporting residue-level evidence, ColBERT-PPI provides a computational basis for investigating physiological and disease-associated interactions and prioritizing proteins and interaction regions for therapeutic exploration. Code is available at https://github.com/UR-Free/ColBERT-PPI.

## Introduction

Interactions among biological macromolecules underpin a wide range of physiological processes, and disruption of these networks contributes to disease development^1,2^. Among these interactions, protein–protein interactions (PPIs) organize networks involved in cellular signaling, gene regulation and macromolecular assembly^3-5^. Understanding these networks entails identifying both the interacting partners and the sites through which they bind. Mapping these sites helps explain how disease-associated variants alter specific interactions^6,7^ and guides the design of binders that engage selected protein surfaces^8,9^. However, experimental maps capture only a fraction of the potential interaction landscape^10^. Computational screening can help prioritize candidate partners, but linking these predictions to the residues that form the binding interface would also help explain how the partners interact. Connecting partner identification with interaction-site evidence is therefore important for investigating molecular function and guiding therapeutic discovery.

Protein language models (PLMs) have become widely used representations for protein structure prediction, functional characterization and protein design^11-14^. For PPI prediction, PLM-based models differ in how the two proteins are encoded and when residue-level information from the two proteins is compared. Cross-encoding approaches such as PLM-interact and PPLM-PPI jointly process protein pairs, allowing information from both proteins to influence their representations, but requiring a new forward pass for each candidate pair^15,16^. Single-vector retrieval approaches such as FlashPPI and RaftPPI instead generate reusable representations for individual proteins before candidate comparison^17,18^; this avoids repeated pairwise encoding but compresses each protein before its interaction partner is known. Because a protein can use different regions of its surface to interact with different partners^19-21^, early compression may discard the local information needed to distinguish one partner from another. The central challenge is therefore to identify candidate partners at scale while retaining the residue-level evidence needed to localise their potential interaction sites.

Late-interaction models from information retrieval offer a way to resolve this tension. ColBERT and ColBERTv2^22,23^ retain contextualized representations of individual tokens and delay fine-grained comparison until a query and candidate are matched. This design makes token representations reusable across candidate comparisons while preserving local information until the matching stage. Translated to PPI screening, this principle could connect partner ranking and site localisation through the same residue-level comparison, while keeping monomer representations reusable across the candidate library.

Guided by this principle, we developed ColBERT-PPI, a structure-aware multi-vector framework that connects protein partner prediction to interaction-site evidence. ColBERT-PPI uses predicted monomer structures^24^ to obtain residue representations from the structure-aware protein language model SaProt^25^. These representations are cached for reuse, and residue compatibility matrices are constructed only after a candidate pair is formed. Calibrated smooth MaxSim aggregates each matrix into a partner-ranking score, while the underlying residue matches remain available for localisation. Partner prediction and the local readout thus arise from the same comparison.

We evaluated ColBERT-PPI on the PINDER labelled test set^26^ and an independent human yeast two-hybrid screen^27^, comparing it with recent PPI models based on reusable single-vector representations or pairwise encoding. ColBERT-PPI outperformed the external comparators on both benchmarks and exceeded its single-vector architecture control in key retrieval metrics. We then tested whether the same comparisons could reveal where interactions occur. Compatibility matrices highlighted experimental interfaces and gave different regional readouts for different partners of the same monomer. Finally, we extended residue-level comparison to RNA partners, finding that PPI-trained protein representations improved protein–RNA retrieval after task-specific fine-tuning. Together, these evaluations connect PPI partner identification to local interaction evidence and test the value of the learned protein features for adaptation to RNA partners.

## Results

### The ColBERT-PPI architecture

ColBERT-PPI is designed to identify candidate protein partners while retaining the residue-level evidence needed to localize their interactions. This requires a representation that preserves local features until a partner is considered. We therefore use SaProt to represent each residue of a predicted monomer through its amino-acid identity and a Foldseek 3Di token describing local geometry^11,24,25,28^. A binary role indicator then maps each monomer into query and candidate contexts within the encoder, generating a separate cache for each role. We cache the resulting residue vectors so that the same protein can be compared with many partners without repeating monomer encoding.

Once a candidate pair is formed, the cached vectors allow us to ask which residues match across the two proteins. Their cosine similarities form a residue compatibility matrix, with each entry describing one possible residue pair. For protein-level retrieval, some residue vectors match many other residues strongly, so a high similarity alone may not indicate a partner-specific match. We account for this by measuring how strongly each residue vector matches a fixed reference set of training residues and using these background similarities to adjust each residue-pair score. This correction follows the rationale of cross-domain similarity local scaling (CSLS), which adjusts vector similarities for neighbourhood similarity on both sides^29^. ColBERT-PPI adapts this correction to individual residues. Rather than averaging similarities to the nearest neighbours, it uses smooth aggregation to estimate each residue’ s background similarity against a fixed set of training residues. These background values remain the same when the candidate library changes, so they can be computed once and stored with the monomer vectors. These background estimates support retrieval calibration, while the raw cosine matrix provides the localization readout (Fig. 1b).

**Fig. 1.**
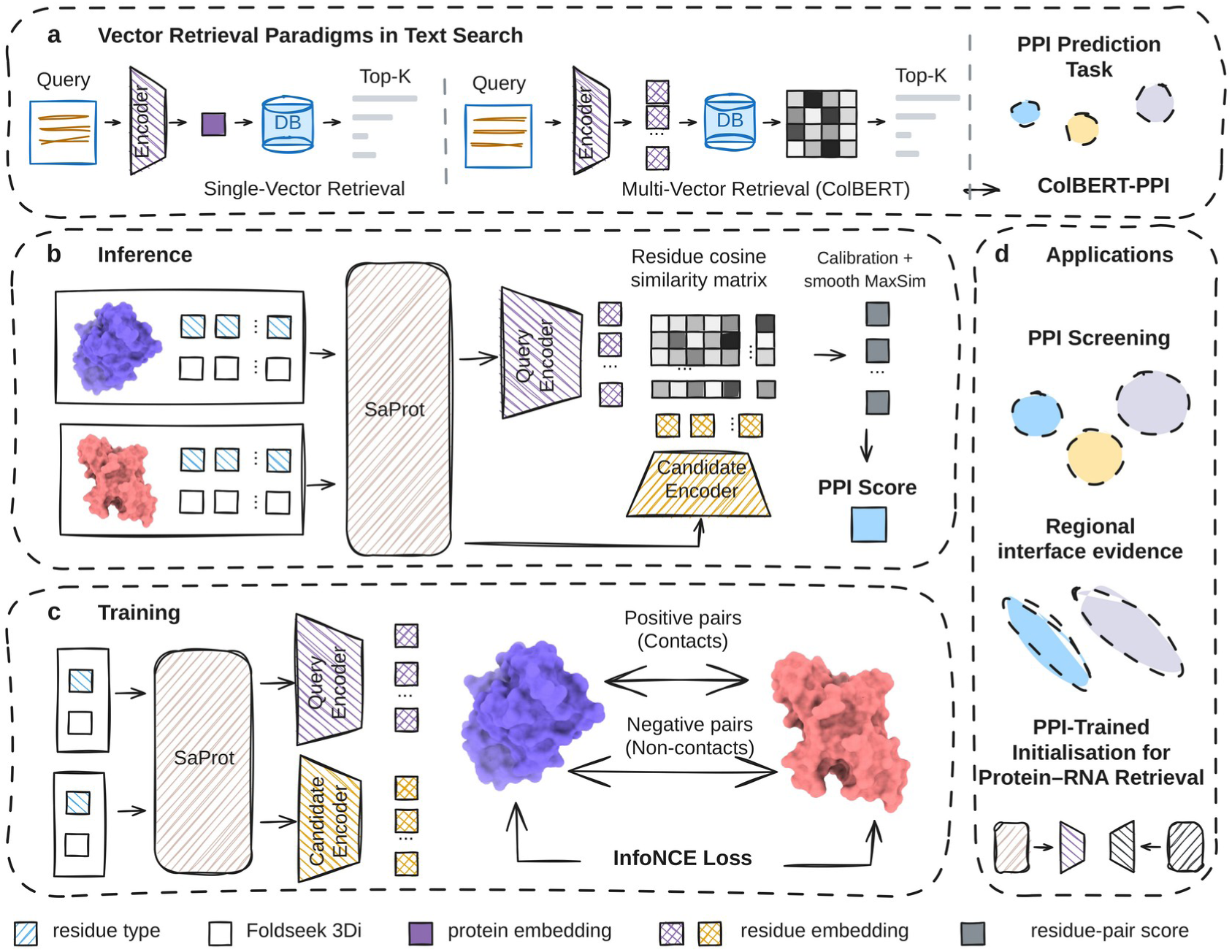
ColBERT-PPI retains partner-dependent residue evidence for retrieval and localization. **a,** Single-vector and multi-vector retrieval paradigms, and their application to protein partner prediction. **b,** Predicted monomers provide amino-acid and 3Di inputs to SaProt and shared role-conditioned context layers, producing reusable query and candidate residue caches. Raw residue cosine similarities provide the localization matrix. For retrieval, training-reference calibration adjusts these similarities and smooth MaxSim aggregates matches in both chain directions; the two explicit encoder role assignments are averaged for the final retrieval score. **c,** Experimental complexes provide contact supervision during training, while predicted monomers supply the inputs. Experimental complexes are not required for inference. **d,** The retained residue evidence supports PPI screening and regional localization, and PPI-trained protein weights initialize subsequent protein–RNA adaptation.

To obtain a protein-level retrieval score, ColBERT-PPI aggregates the calibrated residue compatibility matrix using smooth MaxSim. Classical ColBERT performs this step by keeping the strongest candidate-token match for each query token^22^. Our smooth MaxSim readout gives the strongest matches the most weight while retaining contributions from alternatives. It summarizes the matches available to each residue, then averages over residues in each chain and combines the two directions. This averaging prevents a longer chain from accumulating more evidence simply because it contains more residues. The query and candidate roles introduced during encoding are also exchanged, and the two resulting pair scores are averaged. The raw cosine matrix is retained for localization, linking both readouts to the same learned residue representations.

Once encoding, residue-level comparison and protein-level scoring are defined, training connects the framework to experimental contact geometry while retaining predicted monomers as model inputs (Fig. 1c). After predicted-monomer mapping, the PINDER training set contained 34,017 prepared records^26^. Contacts were defined by Cβ distances below 8 Å (Cα for glycine), while separations above 12 Å or alternative residue pairings supplied negative examples. These labels guided fine-tuning of LoRA adapters in SaProt and the downstream residue-level layers, teaching the representations to distinguish contacting from non-contacting residue pairs. Experimental complexes therefore provide supervision during training, whereas inference uses predicted monomers alone. In this way, the model learns to rank candidate partners using residue-level features shaped by experimentally observed contacts.

### ColBERT-PPI improves protein partner retrieval

Having defined how the framework links pair scores to local evidence, we first evaluated its ability to rank labelled interaction partners within a finite candidate set. For this evaluation, we constructed a labelled test set by combining known interactions from the PINDER test partition with database-filtered operational negatives at a 1:100 ratio. This follows the established PPI evaluation approach of testing whether models can distinguish known interactions from sampled background pairs^17^. We compared ColBERT-PPI with four recent PLM-based PPI models spanning single-vector retrieval and cross-encoding. FlashPPI and RaftPPI independently compress each monomer into a reusable single vector, making them the closest external architectural comparators for ColBERT-PPI’s multi-vector design. All three retrieval models were trained on the same PINDER training set. The pairwise encoders PPLM-PPI and PLM-interact were evaluated using their released task-specific checkpoints and published configurations (Supplementary Table 1). On the PINDER labelled test set^26^, ColBERT-PPI reached an AUPRC of 0.482 versus 0.151 for FlashPPI, the strongest external comparator by this metric (Fig. 2a). Labelled positives were enriched at higher scores relative to operational negatives (Fig. 2b). ColBERT-PPI also achieved the highest mean reciprocal rank and Hit@1, Hit@5 and Hit@10 across the complete candidate universe (Fig. 2c). ColBERT-PPI therefore recovered more labelled partners at the earliest screening ranks than the evaluated external methods.

**Fig. 2.**
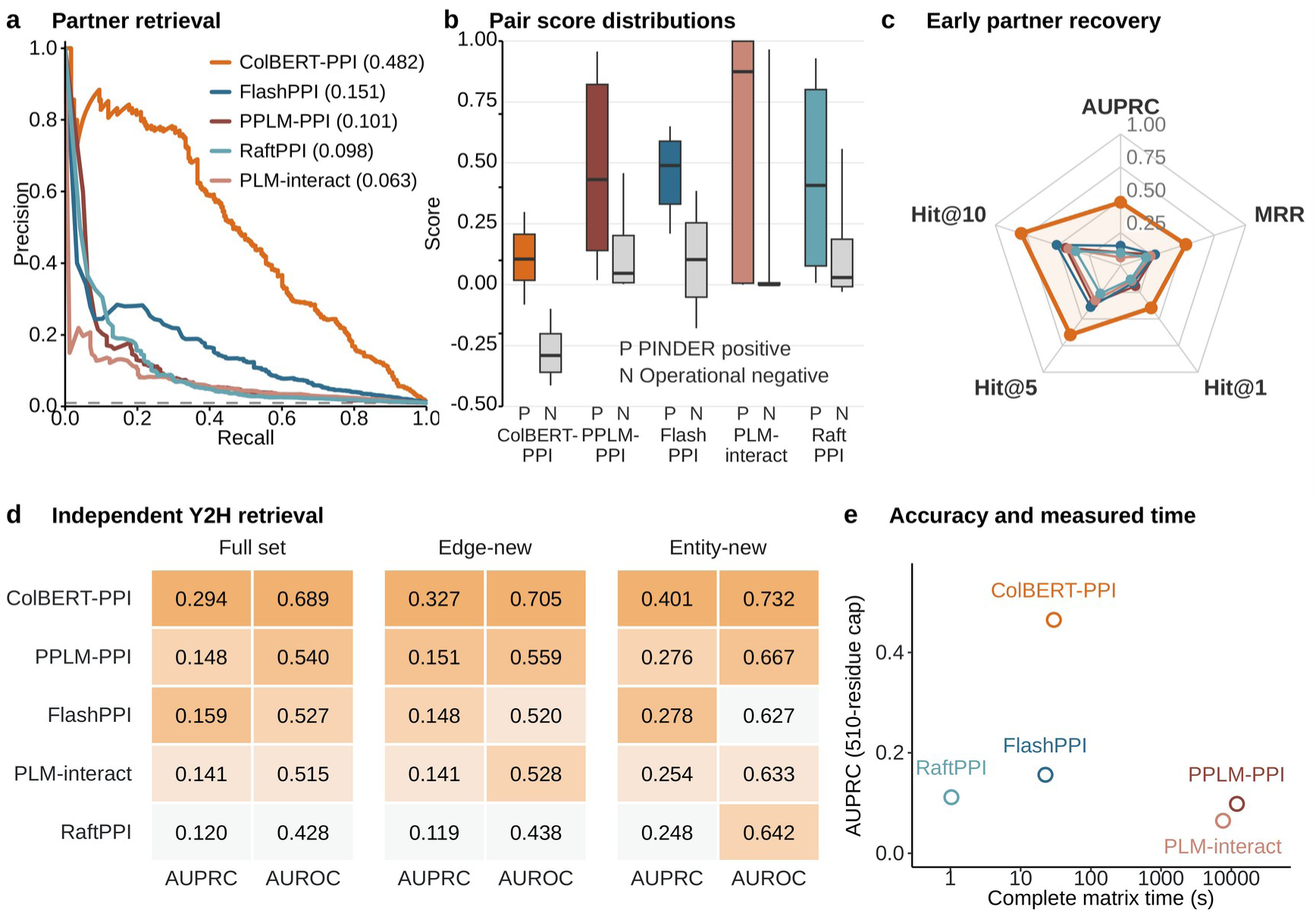
Protein-protein interactions retrieval. **a,** PINDER precision-recall curves on the PINDER labelled test set (243 positives and 24,300 operational negatives); parenthetical values show AUPRC and the dashed line shows prevalence. **b,** Positive and operational-negative score distributions for the same set. The boxes show the interquartile range, center lines show medians and whiskers show the 10th-90th percentiles. **c,** Radar comparison of all five models using PINDER AUPRC, mean reciprocal rank (MRR), Hit@1, Hit@5 and Hit@10 on the original 0-1 scale of each metric; AUPRC uses the labelled test set, whereas MRR and Hit rates use the complete candidate universe. **d,** Y2H AUPRC and AUROC for the full 2,493-pair, edge-new 2,110-pair and entity-new 165-pair sets; darker shading indicates a higher within-set rank. **e,** PINDER AUPRC recalculated on the same 24,543 labelled pairs against accelerator time for the complete 248 × 248 PINDER matrix, using the same 510-residue cap for both AUPRC and timing and one synchronized measurement per method after one warm-up.

To assess generalizability beyond the PINDER benchmark, we applied the same model without any retuning to an independent human yeast two-hybrid screen that uses experimental labels^27^. Among the five evaluated models, ColBERT-PPI achieved an AUPRC of 0.294 and an area under the receiver operating characteristic curve (AUROC) of 0.689 on the full Y2H set, the highest values for both metrics. It also ranked first on both metrics for the edge-new and entity-new subsets, which excluded previously exposed pairs and protein entities, respectively, under the study’s exposure protocol (Fig. 2d). The retrieval advantage thus extended to an independent assay-based benchmark.

The practical value of these retrieval gains also depends on the computational cost of screening candidate partners. Under the common 510-residue cap, ColBERT-PPI achieved the highest AUPRC (0.465) among the evaluated models and completed the full PINDER timing matrix in 29.7 s, placing it at the high-accuracy end of the measured performance-time Pareto frontier (Fig. 2e). Role-specific caches avoided repeated backbone inference, while exact pair scoring constructed a residue compatibility matrix for each protein pair. Computing the per-residue training-reference background was included in encoding time. Phase-specific timing and length-dependent scaling are provided in Supplementary Table 2 and Supplementary Fig. 1. These measurements show that encoding reuse makes the residue comparisons underlying both retrieval and localization practical for the evaluated candidate set.

### ColBERT-PPI links partner retrieval to regional interface evidence

Beyond ranking partners on PINDER and Y2H, we asked whether the underlying residue comparisons could also identify regions involved in binding. We tested whether raw cosine similarities between the learned residue vectors identify experimentally defined interface residues. For each residue, we took the maximum matrix score across the partner chain and evaluated the resulting profiles on a fixed cohort of 226 positive complexes from the PINDER labelled test set. ColBERT-PPI reached a mean interface-residue AUROC of 0.637, compared with 0.568 for FlashPPI and 0.497 for RaftPPI (Fig. 3a). The residue representations underlying partner retrieval therefore also localised interface-enriched regions, without a separately trained interface classifier.

**Fig. 3.**
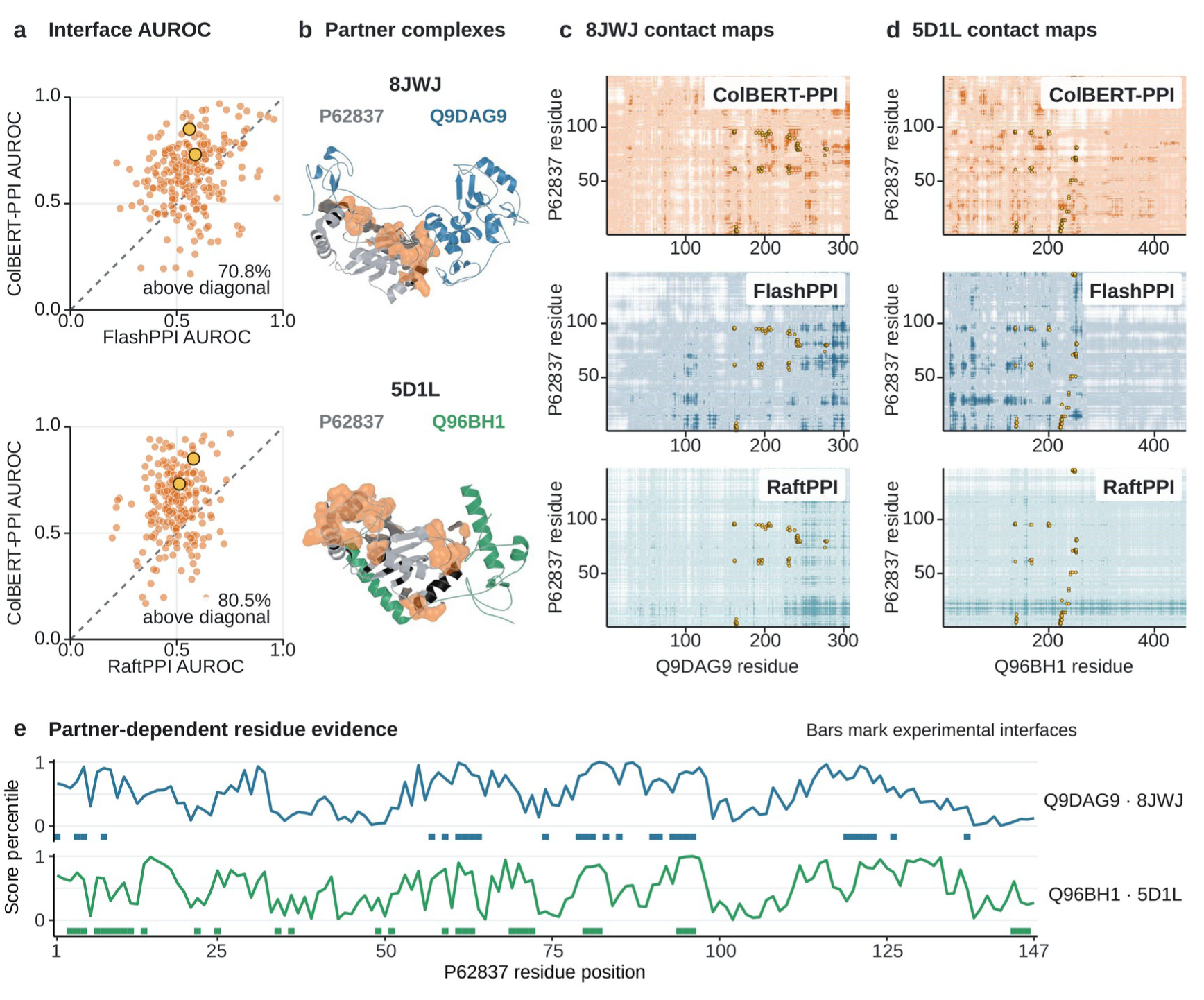
ColBERT-PPI retains regional interface evidence. **a,** Per-complex interface-residue AUROC for ColBERT-PPI against FlashPPI and RaftPPI across the fixed interface evaluation cohort (n = 226). Diagonals denote equal performance, gold points identify the P62837 examples in b-d and labels give the fraction above the diagonal. **b,** Experimental structures 8JWJ and 5D1L, pairing the same complete 147-residue P62837 monomer (light grey) with Q9DAG9 (blue) or Q96BH1 (green); orange surface marks high-scoring P62837 regions and black segments mark the experimental interface. **c-d,** Raw residue cosine matrices for ColBERT-PPI and native residue-pair score matrices for FlashPPI and RaftPPI for 8JWJ (**c**) and 5D1L (**d**). Colors follow Fig. 2, color intensity gives the within-matrix score percentile, and gold-outlined points mark experimental residue contacts below 8 Å. **e,** ColBERT-PPI residue-score profiles obtained by maximum-over-partner projection of the raw cosine matrices for the same P62837 monomer paired with Q9DAG9 (blue) or Q96BH1 (green). Scores are shown as within-profile percentiles across all 147 residue positions. Colored bars below each profile mark annotated experimental interface residues.

Given the sparsity of interface residues, we complemented AUROC with AUPRC to assess the robustness of regional enrichment across metrics^30^. The model ordering was unchanged, with mean AUPRC of 0.362 for ColBERT-PPI, 0.316 for FlashPPI and 0.257 for RaftPPI (Supplementary Fig. 2). Final-layer inter-protein attention from a released PPLM-PPI checkpoint was analysed separately and produced lower cohort means on both interface metrics (Supplementary Fig. 2). The consistent ordering across both metrics supported interface enrichment at the cohort level.

Comparing the same predicted monomer with two different partners then allowed us to test whether its regional evidence changed with partner identity. We selected the example solely by minimizing overlap between the experimental interfaces, without reference to model scores or performance. This procedure identified the 147-residue protein P62837 paired with Q9DAG9 in 8JWJ and Q96BH1 in 5D1L; its two interfaces had a Jaccard index of 0.240 (Figs. 3b-e). The P62837 map generated with Q9DAG9 reached an interface AUROC of 0.828 against the 8JWJ interface, compared with 0.712 for the map generated with Q96BH1. Against the 5D1L interface, the corresponding values for the native and alternative partners were 0.666 and 0.614, respectively, indicating that changing the partner altered the residues contributing to the local evidence. To test how strongly the retrieval evidence was concentrated, we ranked each query’s residues by their contributions to its score with the known interaction partner. We then retained the top 5% and used this fixed residue mask to rescore the candidate library. Across the PINDER systems, this intervention increased AUPRC from 0.482 with complete inputs to 0.642 (Supplementary Fig. 3 and Supplementary Table 3). Focusing on residues selected for the known partner increased its score advantage over competing candidates. This gain suggests that a small subset of residues can retain the evidence needed to distinguish the known partner from competing candidates. Together, the shared-monomer example and score interventions connect the identity of the candidate partner to the local evidence supporting its predicted interaction.

### Multi-vector comparison supports partner retrieval and localisation

ColBERT-PPI recovered more labelled partners than the external comparators, and the same residue comparisons highlighted experimental interface regions. We next asked how multi-vector comparison and 3Di structural context contributed to these two capabilities. The single-vector model retained amino-acid and 3Di inputs but pooled residue representations before comparison and used a protein-level contrastive objective; the sequence-only model retained multi-vector comparison without 3Di. For retrieval, both multi-vector models used the same calibrated smooth MaxSim parameters, with reference banks drawn from their respective training representations, while the single-vector model used pooled cosine similarity. All three models were evaluated on PINDER edge-new and entity-new pairs; the latter tests generalisation when neither protein was seen during training^31^. ColBERT-PPI achieved the highest AUPRC and AUROC in both settings (Figs. 4a, b). Its edge-new AUPRC was 0.482, compared with 0.481 for single-vector and 0.380 for sequence-only. On entity-new pairs, the corresponding values were 0.625, 0.557 and 0.483. Structural input therefore strengthened multi-vector retrieval on PINDER, and the complete model exceeded the single-vector control with both familiar and unseen protein entities. Sequence- and structure-distant analyses provide additional fixed-cohort comparisons (Supplementary Table 4).

**Fig. 4.**
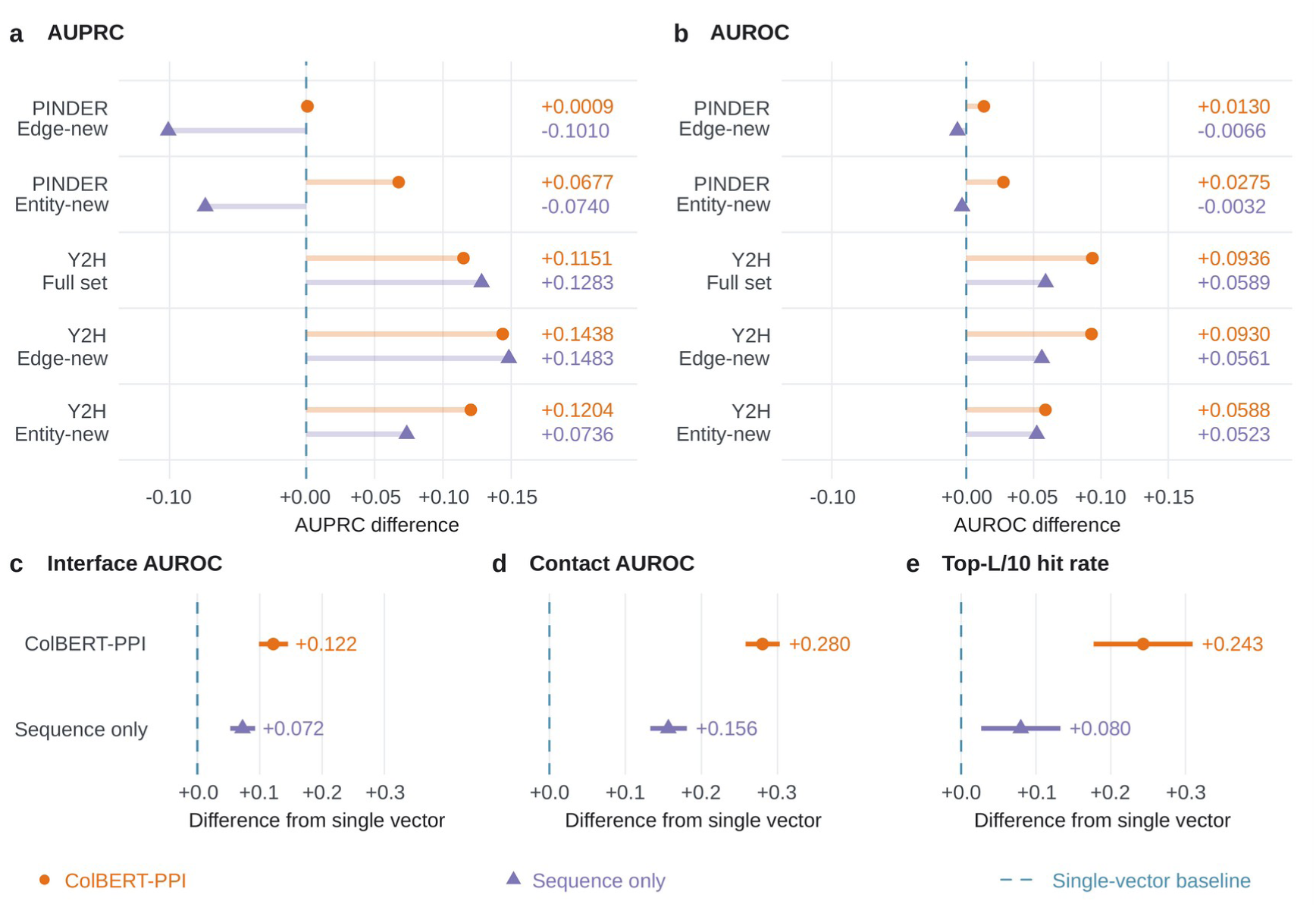
Multi-vector comparison supports retrieval and preserves residue-level evidence. **a-b**, AUPRC (**a**) and AUROC (**b**) differences from the single-vector control on PINDER edge-new and entity-new pairs and the independent Y2H full, edge-new and entity-new sets. Positive values favor the indicated multi-vector model. For retrieval, both multi-vector models use calibrated smooth MaxSim; the single-vector control uses pooled cosine similarity. **c–e**, Mean paired differences in interface AUROC (**c**), contact AUROC (**d**) and the fraction of complexes with at least one experimental contact among the top-L/10 residue pairs (**e**), across 226 complexes. Localization uses raw residue cosine matrices for both multi-vector models, without background correction or smooth aggregation. Dashed zero lines denote the single-vector baseline. Retrieval points show fixed-set differences; horizontal intervals in c– e are 95% percentile confidence intervals from 5,000 paired-complex bootstrap resamples.

The Y2H screen provided a complementary setting in which to assess the contributions of multi-vector comparison and structural context using a distinct source of interaction labels (Figs. 4a, b). On the full set, sequence-only and complete ColBERT-PPI reached AUPRCs of 0.307 and 0.294, respectively, compared with 0.179 for the single-vector control. The same ordering held for edge-new pairs (0.331, 0.327 and 0.183). On entity-new pairs, the complete model led with 0.401, followed by sequence-only at 0.354 and single-vector at 0.280. Both multi-vector models also exceeded single-vector AUROC in all three sets, with the complete model giving the highest AUROC. The higher sequence-only AUPRC on the full and edge-new sets shows that the advantage of retaining residue-level comparisons extends to sequence-based retrieval, while the complementary PINDER results identify a setting in which structural context further improves ranking.

Whereas retrieval metrics establish which partners are recovered, regional readouts reveal where the supporting evidence is localized. Using the raw residue cosine matrices, ColBERT-PPI achieved a mean interface AUROC of 0.637 and contact AUROC of 0.806, compared with 0.515 and 0.526 for the single-vector model (Figs. 4c, d). Sequence-only multi-vector comparison reached 0.588 and 0.683, retaining contact-level information while structural input strengthened localization. ColBERT-PPI recovered at least one experimental contact among the top-L/10 residue pairs in 31.9% of the 226 complexes, compared with 15.5% for sequence-only and 7.5% for single-vector (Fig. 4e). The ablations therefore support the use of multi-vector representations for both partner retrieval and localization, with 3Di input providing additional regional and contact-level evidence.

### PPI training improves protein-RNA retrieval after adaptation

Transfer learning suggests that related tasks and domains can benefit from shared representations^32^. ColBERT-PPI’s late-interaction architecture provides a natural route to such transfer because it encodes each partner separately before comparing their local representations. This separation allows the protein branch to be reused alongside an encoder for a different molecular partner, extending the framework beyond PPIs to protein–partner retrieval more broadly. We examined this possibility in protein–RNA retrieval, building on advances in modelling RNA and its interactions with proteins^33-36^. We therefore hypothesized that protein features learned through PPI contact supervision could help identify RNA partners after adaptation. To test this, we initialized the protein branch with PPI-trained weights and fine-tuned the model on protein–RNA complexes from BioLiP2. BioLiP2 curates biologically relevant protein –ligand interactions from experimentally determined structures in the Protein Data Bank, including protein–RNA complexes with annotated binding sites^37^. We used residue–nucleotide contacts from these complexes to learn compatibility with RNA partners within the same multi-vector comparison framework. The RNA branch used the pretrained ERNIE-RNA language model^38^, with nucleotide sequences as the only RNA input and no experimentally determined or predicted RNA structures. Sequence-cluster components defined disjoint partitions, and predicted-monomer mapping yielded 9,392 training records. Following the same test-set construction principle as PINDER, we paired structural positives with database-filtered, taxonomy-stratified operational negatives. All models used identical evaluation labels. We compared released Graph-RPI and Transfer-RPI checkpoints with same-split RPISeq-RF and SeqMG-RPI controls^39-42^.

Within our model family, we compared multi-vector and single-vector scoring with either SaProt-only initialization (denoted de novo) or PPI initialization. All conditions were evaluated on the same database-filtered benchmark after exact source-sequence exclusion. PPI-initialized multi-vector scoring achieved the highest mean exact-entity AUPRC, reaching 0.0201 compared with 0.0155 for the PPI-initialized single-vector control, a relative improvement of 30.2% (Fig. 5b). Under SaProt-only initialization, the multi-vector and single-vector models reached AUPRCs of 0.0148 and 0.0156, respectively. PPI initialization increased mean multi-vector AUPRC by 35.7%, with improvements in all three seeds. Single-vector mean AUPRC decreased by 0.8%. Even without PPI initialization, multi-vector retrieval exceeded all four external comparators, whose highest AUPRC was 0.0106. These comparisons support the combination of residue-level scoring and PPI-trained initialization for RNA partner retrieval.

**Fig. 5.**
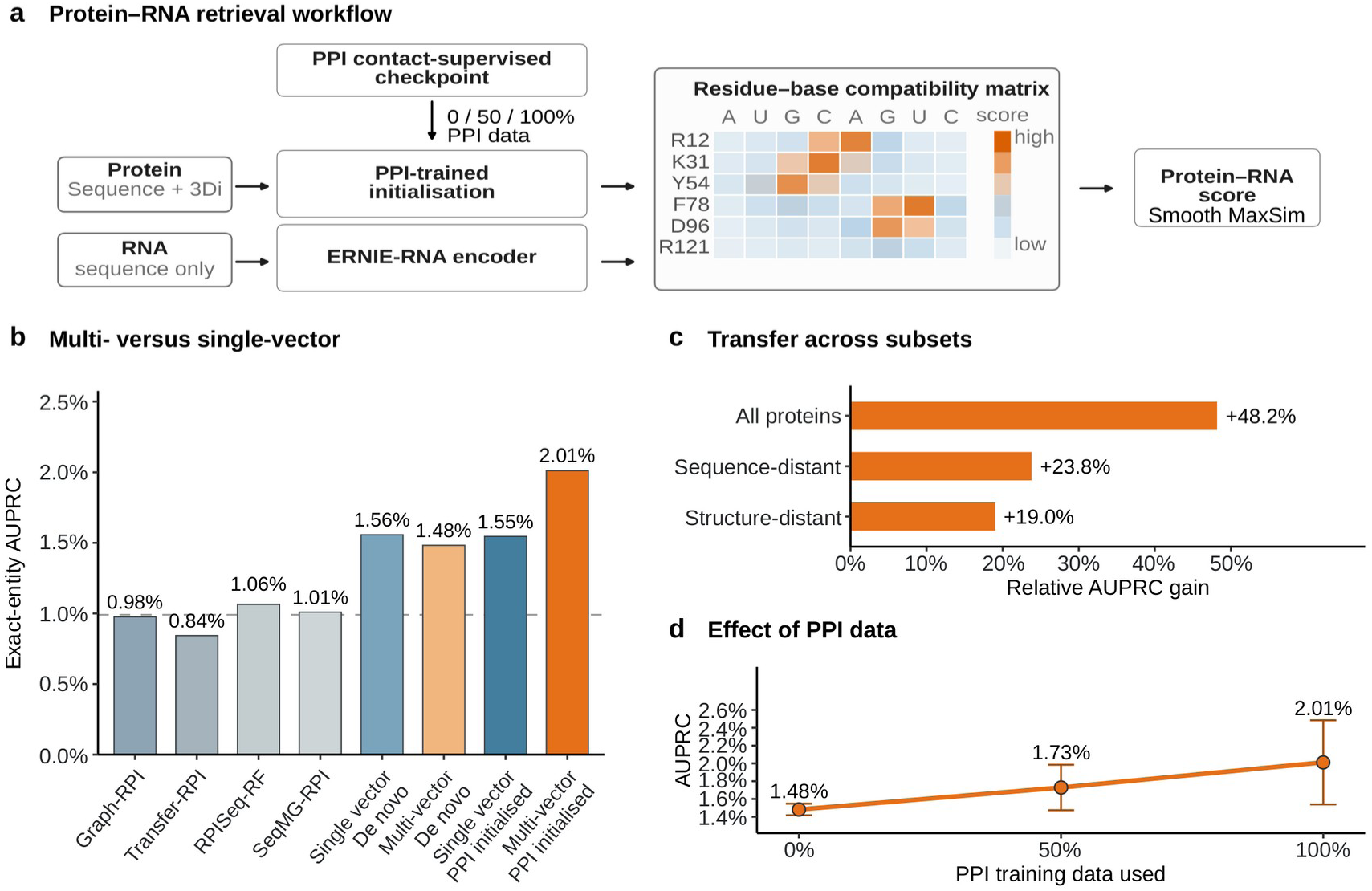
PPI initialization strengthens multi-vector protein-RNA retrieval after fine-tuning. **a,** PPI contact-supervised initialization before protein-RNA fine-tuning and residue-base scoring. **b,** Exact-entity AUPRC after exact source-sequence exclusion, using 576 structural positives and 57,600 database-filtered operational negatives from a candidate grid of 447 proteins and 187 RNAs; dashed line, positive prevalence. SaProt-only (labelled de novo) and PPI-initialized conditions compare multi-vector and single-vector models. Internal bars are three-run means. **c,** Relative gains in mean multi-vector AUPRC from PPI initialization on the full, sequence-distant and structure-distant protein cohorts (459, 189 and 94 proteins). Cohorts were defined by sequence and structural exclusions relative to PPI training (Methods); labels show gains over SaProt-only initialization. Each cohort used identical labels across methods. Database-filtered, taxonomy-prioritized negative sampling targeted 1 to 100, with 1 to 93.3 for structure-distant because fewer eligible negatives were available. **d**, Effect of PPI-data fraction on mean multi-vector AUPRC in the same labelled set as b. The 0% and 100% endpoints match b; error bars show s.d. across three seeds.

We next varied the amount of PPI training data used for protein-side initialization while keeping RNA initialization and downstream training unchanged. On the same evaluation set used for the model comparison, after exact source-sequence exclusion, mean exact-entity AUPRC was 0.0148 with SaProt-only initialization and 0.0173 and 0.0201 after initialization with 50% and 100% of the PPI data, respectively (Fig. 5d). The ordered increase suggests that exposure to more PPI training data provides a progressively more useful starting point for learning RNA partner compatibility.

Finally, we used nested exclusion filters to assess the transfer benefit beyond close sequence and structural analogues in the PPI training data. Within each set, positives and operational negatives were selected using the same rules for both initialization conditions. Relative to SaProt-only initialization, full PPI initialization changed mean multi-vector AUPRC by +48.2% across all proteins, +23.8% on the sequence-distant set and +19.0% on the structure-distant set (Fig. 5c). The average transfer benefit therefore persisted across all three sets, including proteins lacking close sequence and structural matches in the PPI training data. This transfer pattern extends the value of PPI contact-supervised protein representations to RNA partner retrieval after task-specific adaptation.

## Discussion

PPI prediction is important for understanding cellular processes and disease mechanisms, yet identifying interaction partners from a large candidate space remains challenging. Many existing approaches condense each protein into a single vector before partner comparison, which facilitates efficient retrieval but can obscure the local residue-level information that distinguishes interactions with different partners. In this study, we developed ColBERT-PPI, to our knowledge the first multi-vector framework designed for protein partner retrieval, which retains residue-level protein representations until candidate partners are compared. This design connects partner ranking to the residue-level matches supporting each prediction, extending the comparison beyond the pooled representations or pairwise encoding used by existing PPI approaches^15-18,43-46^. This partner-dependent comparison is particularly relevant when a protein is evaluated against multiple partner candidates, because different comparisons can draw on different regions. ColBERT-PPI showed a larger retrieval advantage on PINDER than on Y2H, two benchmarks that assess partner prediction in different settings. PINDER explicitly evaluates each query against multiple candidate partners, whereas Y2H mainly comprises independently assayed pairs. The former therefore tests partner discrimination for a shared query more directly. As practical partner screening more closely resembles the PINDER scenario, ColBERT-PPI’s stronger performance on this benchmark highlights its potential for real-world retrieval.

The protein-RNA experiments support combining multi-vector comparison with PPI-trained protein representations. After adaptation, this combination achieved the highest mean AUPRC on the common database-filtered benchmark, and increasing the amount of PPI training data produced an ordered improvement in mean retrieval performance. Starting from similar retrieval performance under SaProt-only initialization, the multi-vector model gained more from PPI training, linking its advantage to the transfer of interaction-informed protein features. PPI initialization also improved mean retrieval performance on both sequence- and structure-distant protein cohorts. These findings support contact-supervised protein features as a useful starting point for learning interactions with RNA. The PPI ablations clarify the contribution of structural information within this framework. Both the complete and sequence-only multi-vector models exceeded the single-vector control on Y2H, with sequence-only inputs yielding the highest AUPRC on the full and edge-new sets. The complete model performed better on PINDER and provided stronger localization, indicating that 3Di contributes differently across tasks. Together, these results support retaining local representations until partner comparison and adapting interaction-informed protein features through task-specific supervision.

Beyond improving partner retrieval, ColBERT-PPI also reveals local residue matches specific to the candidate partner. The raw cosine matrix provides this regional readout, while background calibration and smooth MaxSim convert the residue comparisons into a retrieval score. This capability distinguishes the framework from single-vector approaches such as FlashPPI and RaftPPI^12,13^, which pool residue information before the partner is known. For studies of physiological and disease-associated interactions, these local readouts could help prioritize regions for mutational testing and focused investigation of interaction sites. However, an inspectable regional readout does not by itself establish a physical binding mechanism. Its correspondence with experimental contact maps remains incomplete, potentially because local residue-pair matching does not explicitly enforce global three-dimensional consistency. Adding geometric or physical constraints, or applying structure-aware refinement to top-ranked pairs, could improve the correspondence between regional evidence and physical contacts. Such refinement would strengthen the use of these predictions in studies of interaction mechanisms and potential therapeutic intervention sites.

Overall, ColBERT-PPI establishes multi-vector comparison as a unifying framework for protein partner retrieval and local interaction evidence. Reusable residue representations allow a protein to be assessed against different partners while preserving the local matches that support each prediction. Their successful adaptation to protein–RNA retrieval further demonstrates the value of PPI-trained protein features beyond the original task. This framework provides a foundation for investigating physiological and disease-associated interactions and prioritizing candidate partners and protein regions for experimental and therapeutic exploration.

## Methods

### Datasets and study design

We evaluated partner ranking on PINDER and human Y2H, assessed regional interface evidence, and tested protein-RNA retrieval. After predicted-monomer mapping, 34,017 records formed the PINDER training set, and validation and test contained 257 and 248 records, corresponding to 253 and 243 canonical-accession interactions^26^. Predicted monomers provided model inputs, while experimental complexes supplied training and localization labels. Structural retrieval tests for PPI and protein–RNA interactions shared a score-independent construction principle, retaining structural positives and sampling database-filtered operational negatives through taxonomy-based tiers towards a target of 100 negatives per positive. Y2H retained its experimental assay labels. We used the published PINDER partition assignment. An independent DataSAIL addendum reported lower scaled leakage for the November 2023 PINDER split than for its interaction-weighted DataSAIL comparison^47^; this published comparison was not a new split analysis performed here.

The full set includes every scored member, edge-new excludes development pairs and entity-new requires both endpoints to be unseen. Sequence- and structure-distant PPI filters were applied separately, as defined under ablation evaluation. Protein–RNA analyses used the nested component-level controls described below.

### PPI structure mapping and residue inputs

PINDER chains were mapped to canonical UniProt accessions and AlphaFold Protein Structure Database monomers or, when unavailable, ESM Metagenomic Atlas structures^11,24^. Coordinates were parsed without docking or complex prediction, and Foldseek generated 3Di sequences^28^.

Experimental chains were matched to predicted monomers by exact substring or local alignment requiring at least 80% coverage and 70% identity, with isoform- and SIFTS-aware remapping below that coverage. Contact pairs had Cβ distances below 8 Å, using Cα for glycine; distant pairs exceeded 12 Å, with the intervening margin excluded.

SaProt-650M combined amino-acid and 3Di tokens^25,28^. Prespecified windows were contact-enriched training windows of at most 384 residues, deterministic N-terminal ranking windows of at most 1,024 residues and construct-aligned Fig. 3 windows of at most 512 residues. Only training-window selection used contact labels.

### ColBERT-PPI architecture

ColBERT-PPI used SaProt-650M-PDB as its shared backbone. LoRA adapters were fine-tuned in the self-attention projections while the remaining backbone weights were frozen^48^. SaProt produced 1,280-dimensional residue vectors. A binary query or candidate role indicator preceded a shared three-layer Transformer with FlashAttention, producing 512-dimensional L2-normalised vectors.^49^ Each monomer was encoded once per role through shared parameters. A shared linear head produced sigmoid residue weights applied once to each pairwise similarity during contact-supervised training. The retrieval readout used the L2-normalised residue vectors without an additional residue-weight or fixed-logit-scale multiplier.

### PPI training objective and optimisation

Training sampled contact-anchored monomer windows and included contacting, interface and distant residues from each complex. For P contacts, xᵢ and yᵢ denote L2-normalised embeddings and aᵢ and bᵢ their sigmoid weights. Equations (1)-(3) define the weighted logits, row- and column-wise log-probabilities and symmetric contact loss, respectively.

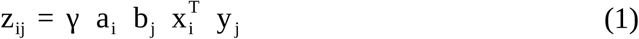

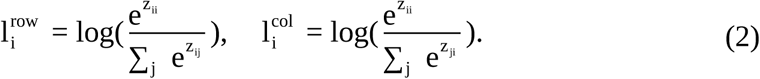

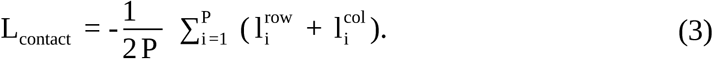

Only contact rows and columns were anchors, and denominators included the P + D sampled residues from the opposite chain. Off-target P × P, P × D and D × P entries were negatives, whereas D × D was excluded. The logit scale was fixed at γ = 1/0.07, with each residue weight entering once. The full objective added 0.01 R to the contact loss, where R = mean[a(1 − a)] + mean[b(1 − b)] over valid residues of the two branches, excluding special tokens and padding. This term favors residue weights near zero or one. No protein-level or AUC-surrogate loss was used.

Trainable components were optimized with AdamW for 80 epochs, with a checkpoint saved after each epoch. Checkpoint selection used separate offline validation runs after training; test data were not used in optimization or selection.

### Late-interaction scoring and model selection

For protein-level retrieval, reference calibration adjusted residue similarities before aggregation using

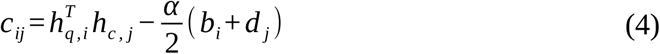

In Equation (4), the background terms bi and dj apply Lτ to cosine similarities between each L2-normalised residue vector and the opposite-role reference bank. Setting α = 0 recovers uncorrected cosine similarities. Smooth MaxSim averages the row-wise and column-wise Lτ summaries of C, approaching maximum matching as τ approaches zero. Pair scores average two independently computed encoder-role assignments. Localization uses the raw cosine matrices as described below.

Each frozen, checkpoint-specific reference bank contained 16 evenly spaced residues from each of 512 deterministically selected training token-sequence identities, giving 8,192 vectors per role. No test data entered the banks. Unlike classical CSLS^29^, we use smooth training-reference summaries instead of nearest-neighbor means and subtract half the combined background at α = 1, retaining the cosine scale. Calibration precedes pooling and preserves cached encoding and local readouts without whitening or weight updates.

Checkpoint selection used fixed smooth MaxSim settings before the final parameter search. For both PPI multi-vector models, all 80 checkpoints were compared by canonical-entity PINDER validation AUPRC at α = 1 and τ = 0.03, using unweighted L2-normalised vectors and checkpoint-specific training-reference banks. No additional residue-weight or fixed-logit-scale multiplier was applied. This selected complete-model epoch 69 and sequence-only epoch 50, with the earliest epoch used to resolve ties. The single-vector model retained its validation-selected epoch 74 and native pooled score.

We then held the selected checkpoints fixed and compared α ∈ {0, 1} and τ ∈ {0.001, 0.003, 0.01, 0.03, 0.1, 0.3}, maximizing complete-model PPI validation AUPRC or mean validation AUPRC across nine multi-vector protein–RNA runs. Each α family used its best validation temperature, with ties favoring the smaller temperature. At these temperatures, α = 1 was enabled when its paired mean log-AUPRC gain exceeded one bootstrap standard deviation across 1,000 entity resamples. Protein endpoints were resampled jointly for PPI and separately from RNA endpoints for protein–RNA, with weights shared across runs. This regularization heuristic selected α = 1 and τ = 0.03 for both PPI multi-vector models and Y2H, and α = 0 and τ = 0.001 for all nine protein– RNA models. Checkpoints and final parameters were fixed before test evaluation; test metrics did not enter either selection step.

### PINDER ranking evaluation

PINDER positives were test-partition interactions absent from earlier partitions^26^. Following the shared structural-retrieval evaluation principle, PINDER operational negatives were selected through a score-independent database-absence hierarchy using PINDER and STRING^50^. These labels are not verified non-interactions. After canonical collapse, validation contained 253 positives and 25,299 operational negatives, while test contained 243 positives and 24,300 operational negatives. The 243 query and 244 candidate entities formed 59,292 possible ordered cells. Eligibility excluded 10 cells overlapping training or validation interactions and a further 1,688 non-positive cells without complete endpoint taxonomy, leaving 57,594 candidate pairs. Structural positives remained eligible regardless of taxonomy. AUPRC and AUROC used the labelled set; mean reciprocal rank and Hit rates included unlabeled cells in the complete eligible universe.

Operational negatives were selected through five ordered tiers of decreasing database coverage. The first retained all fully STRING-mapped, association-free pairs from the strict protocol, including same-species pairs and cross-species pairs whose species combination occurred among positives in that partition. The second admitted fully mapped cross-species pairs without that species-combination match. The remaining three tiers admitted incompletely mapped pairs in the order of species-combination-matched cross-species, same-species and unmatched cross-species pairs. Every tier excluded PINDER interactions and development-edge overlap; mapped STRING associations were excluded.

Deterministic sampling fixed the final negative set without reference to model predictions. Tiers were exhausted in order until there were 100 negatives per positive. A deterministic ordering resolved partially used tiers. Positive labels took precedence, and eligible cells outside the labelled set remained unlabeled ranking candidates.

### Comparator models

ColBERT-PPI, FlashPPI and RaftPPI used the same PINDER training set and matched batch sizes while retaining their native objectives. ColBERT-PPI used epoch 69 selected as described above; validation selected FlashPPI epoch 27 and RaftPPI step 670. FlashPPI used gLM2-650M-UR50 with protein-pair contrastive and contact objectives, whereas RaftPPI used ESM2-8M fine-tuning and rank-1 attention^11,18^.

PPLM-PPI used its released paired 650M encoder and pooled probability ensemble with a 1,022-residue cap. PLM-interact used its released ESM2-650M human checkpoint to jointly encode up to 1,603 concatenated tokens and return a CLS-based probability. The separate PPLM-Contact architecture was not evaluated. Neither comparator was retrained on PINDER.

### Scoring conventions and computational timing

Duplicate PINDER pairs used maximum score and logical-OR labels, with positives taking precedence. ColBERT-PPI, PPLM-PPI and PLM-interact scores were order-averaged; RaftPPI was order equivalent. Fig. 2b showed the unmodified calibrated smooth MaxSim score for ColBERT-PPI and removed the fixed FlashPPI temperature scale; other outputs were unmodified. SaProt and both role-conditioned ColBERT-PPI outputs were cached before pairing. For N proteins, joint encoding requires NZ backbone evaluations versus 2N role-specific encodings, while exact scoring remains O(NZLZd). Timing therefore tests encoding reuse, not subquadratic retrieval.

Timing covered the complete 61,504-cell PINDER matrix after warm-up on identical A100 GPUs. The matrix compared 248 query records with 248 candidate records. Each run used one method-specific warm-up and one synchronized measurement on an A100 GPU. Inputs had a 510-residue N-terminal cap, with float32 computation and bidirectional averaging where applicable. Timing included tokenization, device transfer, forward passes, aggregation and host transfer, but excluded model loading, metrics, serialization and offline reference-bank construction. Per-residue reference comparisons were included in encoding, and their cached scalars contributed to cache size. Total time used an outer timer, so it can differ slightly from the sum of separately timed phases. Fig. 2e computed AUPRC from these same capped score matrices over the 24,543 labelled pairs; Figs. 2a-c instead used the main 1,024-residue cap.

Scaling experiments distinguished a growing all-pairs library from repeated screening with one query. Supplementary Figs. 1a, c evaluated nested all-pairs libraries of up to 248 proteins under a 510-residue cap, encoding every distinct input in both roles. Panel b varied sequence length in a fixed 32 × 32 matrix. Panel d screened one fixed query against increasing numbers of candidates, up to 248. Its cached condition encoded each distinct monomer once per role, whereas its re-encoding condition repeated those encodings for every pair. Both included encoding, cache materialization and scoring. Thus, the largest workloads in a and d contained 61,504 and 248 pairs, respectively, and their total times are not directly interchangeable. Each scaling point used three synchronized repeats, reported as the median and range.

### Independent Y2H evaluation and exposure control

We used published human Y2H-v1 measurements to evaluate protein partner retrievalZZ. The tested candidates originated from an AlphaFold/RoseTTAFold-predicted human interactome. Of 3,222 candidate pairs, 2,676 had usable experimental labels, comprising 376 detections and 2,300 non-detections. Restricting the analysis to pairs scored by all methods yielded 2,493 pairs, including 369 detections. Of 183 exclusions, 180 exceeded length eligibility, one failed sequence mapping and two lacked ColBERT structural features. The PPI scoring parameters were applied unchanged to Y2H. Potential PPLM-PPI training exposure was mapped against 474,085 human supervised records and 1,681 human STRING pretraining pairs, resolving 14,983 UniProt entities. Y2H edge-new removed edges found in either the mapped PPLM-PPI exposure or ColBERT-PPI training set; entity-new excluded proteins found in either set. These rules formed the prespecified exposure protocol for this study and preceded metric calculation. The protocol does not exhaust every possible species, PDB, homologue or other pretraining exposure of external models.

### Interface readout and shared-monomer case selection

The interface evaluation cohort comprised 226 PINDER labelled-test positives with aligned monomers, retained contacts and both interface metrics defined; two mapped complexes without retained contacts were excluded. The ranking input contained 248 complex records corresponding to 243 canonical interactions. The shared AFDB-monomer window inventory covered 228 of these interactions; the other 15 canonical interactions were outside this fixed localization inventory. Two of the 228 complexes had no retained contacts after projection, leaving 226 with both interface metrics defined. Localization windows were capped at 512 residues per chain. Native-matrix evaluation required a paper-defined residue-pair output without a newly trained interpreter; Fig. 3 therefore compared the three eligible PINDER-trained models. For ColBERT-PPI, each matrix entry was the cosine similarity between two L2-normalised output residue vectors. We averaged the forward raw cosine matrix with the transposed, independently computed reverse-role raw cosine matrix. No background correction, smooth aggregation, whitening or additional residue weighting was applied to these localization matrices. FlashPPI averaged bidirectional contact-head logits, and RaftPPI used its rank-1 weighted Gaussian contribution matrix.

Released PPLM-PPI final-layer inter-protein attention, averaged over heads, cross-chain directions and sequence orders, was analyzed separately. PLM-interact was excluded because its published head returns only a pair-level CLS probability. The bare RaftPPI kernel provided a sensitivity control (Supplementary Fig. 2). Residues were scored by the maximum across the partner chain, and per-protein AUROC and AUPRC were averaged within complexes using experimental contacts as labels. Contact AUROC ranked matrix scores for explicitly labelled contact and distant residue pairs within each complex, with mapped contacts below 8 Å as positives and mapped distant pairs above 12 Å as negatives. The top-L/10 hit rate was the fraction of complexes with at least one experimental contact among the top L/10 residue pairs in the full matrix, where L was the shorter evaluated protein length and at least one pair was retained. No interface classifier was fitted.

The shared-monomer illustration used minimum experimental-interface overlap to select P62837, paired with Q9DAG9 in 8JWJ and Q96BH1 in 5D1L (Jaccard 0.240). We identified proteins represented by the same complete predicted-monomer input in at least two complexes and minimized pairwise Jaccard overlap between their experimental interfaces. Selection did not use model scores or performance. For display, partner residues occupied the horizontal axis and P62837 residues the vertical axis, with color intensity giving the within-matrix score percentile and contacts below 8 Å overlaid. Maximum-over-partner projection produced a P62837 residue profile for each partner. Each profile was evaluated against both experimental interfaces on the aligned support of the target complex using AUROC and AUPRC.

### Ablation evaluation and score decomposition

The single-vector ablation kept amino-acid and 3Di inputs but replaced residue comparison with residue-weighted mean pooling, L2 normalization and bidirectional protein InfoNCE. Its ancillary pre-pooling matrix was used only for localization and did not contribute to retrieval scores. Sequence-only retained residue comparison and removed 3Di. Both used the PINDER training set and shared backbone family, context module, training duration and batch size. Validation selected single-vector epoch 74 using its pooled score and sequence-only epoch 50 using the same calibrated smooth MaxSim criterion as the complete model. Final protein-level retrieval applied α = 1 and τ = 0.03 to both multi-vector models with their own training-reference banks; the single-vector model used its temperature-scaled pooled cosine score. The two multi-vector models therefore shared the checkpoint-selection criterion and final retrieval parameters. For localization, both used raw residue cosine matrices with the same averaging of encoder-role assignments; the single-vector model retained its ancillary pre-pooling readout.

PINDER ablation evaluation used edge-new and entity-new, with the full set identical to the edge-new set because all test edges were absent from development. Y2H used full, edge-new and entity-new. Each fixed set used every labelled pair and identical labels across the three models. Fig. 4 reports AUPRC and AUROC differences from the single-vector control for these fixed sets, while Supplementary Tables 4 and 5 report the separate sequence- and structure-distant analyses for ablations and external methods, respectively. All differences were calculated from unrounded estimates as the multi-vector value minus the single-vector value. For localization, 95% percentile confidence intervals for the mean paired differences were obtained from 5,000 bootstrap resamples of the same 226 complexes, using identical sampled complex indices across methods. These intervals quantify complex-sampling uncertainty.

Sequence-distant proteins shared below 40% sequence identity with every PINDER training or validation protein, with at least 80% coverage of both proteins. Structure-distant proteins lacked a Foldseek neighbor meeting at least 80% bidirectional coverage and query and target TM-scores of at least 0.6. These filters were applied separately and differ from edge-new and entity-new exposure scopes. PINDER sequence-distant and structure-distant sets contained 4,601 and 2,964 pairs with 95 and 57 positives. Y2H contained 587 and 522 pairs with 121 and 122 assay detections, respectively.

Score interventions tested whether retrieval evidence was concentrated in partner-dependent residue subsets. Residues were ranked by summed compatibility-times-gradient contributions to calibrated smooth MaxSim, averaged across encoder roles, with residue index resolving ties. Guided removal affected the top 1%, 5%, 10% or 20%, and retention kept the top 5%, 10% or 20%. Each mask was compared with 20 reproducible random masks matched for operation and residue count^51,52^. Masks were applied to cached epoch-69 embeddings without refitting background terms. The native-partner mask was fixed before rescoring the complete candidate library, and the full-input AUPRC was 0.481726.

Competitive filtering compared how the same mask affected the observed partner and its strongest retrieval competitor. For each PINDER system, the strongest non-positive competitor was identified separately in each direction from full-input scores and fixed before intervention. The observed-partner top-5% retention mask was then applied to both comparisons. Score reductions were averaged across directions, with 95% intervals from 5,000 protein-endpoint bootstrap resamples. Negative reduction denotes a score increase after retention. These interventions measure internal score concentration conditional on the observed partner; masked scores are not deployed retrieval estimates or evidence of verified biochemical specificity.

### Protein-RNA data, splits and exposure control

BioLiP2 provided protein-RNA complexes with proteins of 30-512 residues and RNAs of 15-512 nucleotides^37^. Parsing and contact labelling retained 12,584 of 12,974 candidates. Protein and RNA sequences were clustered at 40% and 80% identity, respectively, with 80% bidirectional coverage; connected protein–RNA components defined the partitions. The initial allocation contained 10,229 training, 1,202 validation and 1,153 test records. Predicted-monomer mapping removed 837 training records, leaving 9,392.

Evaluation filtering also excluded sequence clusters used by an earlier protein–RNA training split. Applying the same protein and RNA thresholds removed 335 initial validation and 275 initial test records, leaving 867 and 878. AFDB mapping removed a further 249 and 106 records, respectively. The remaining 1,390 evaluation records were pooled for component-level allocation. A 165-record component containing RNAs longer than 384 nucleotides was reserved as a long-RNA holdout outside Fig. 5. Metadata-balanced allocation of the other components produced 613 validation and 612 test records, with no exact-sequence, cluster or component overlap with training or between evaluation partitions.

PPI-exposure filters defined the nested transfer-evaluation cohorts. Exact accession, protein sequence, SaProt-input and source-structure matches to PINDER training were excluded at the component level. Sequence-distant additionally required every protein to lack a PINDER match at least 40% identity and 80% query coverage. Structure-distant was nested within sequence-distant and required structural evaluability and no Foldseek neighbor at least 80% bidirectional coverage and query and target TM-scores of at least 0.5. Identical protein and RNA inputs defined one entity edge.

### Protein-RNA model, training and evaluation

The protein branch used SaProt amino-acid and 3Di tokens and the RNA branch ERNIE-RNA sequence representations^38^. Separate three-layer context encoders projected both to 512-dimensional L2-normalised vectors before residue-level comparison. No RNA structure was used.

Multi-vector protein encoders were initialized from SaProt alone or from PPI models trained with 50% or 100% of the PINDER training data. The two PPI conditions used matched optimization steps. Protein-side LoRA, context and attention components were transferred and fine-tuned on protein-RNA data. All conditions used the same RNA initialization. The single-vector control used the same model families and BioLiP2 partitions, with residue-weighted pooling, L2 normalization and temperature-scaled cosine scoring. It was initialized from SaProt alone or a validation-selected single-vector PPI model.

Multi-vector training sampled contact and distant residue-nucleotide pairs and used symmetric row/column contrastive loss with attention-weighted logits and off-target negatives. This supervision jointly adapted the transferred protein-side and trainable RNA-side components. Single-vector training used symmetric in-batch contrastive loss over pooled vectors. All conditions were trained with AdamW for 50 epochs using three random seeds and a matched effective batch size of 192. A checkpoint was saved after each epoch for offline validation.

Downstream checkpoint selection used smooth MaxSim before the shared aggregation parameters were tuned. For each of the nine runs at 0%, 50% and 100% PPI-data initialization and three seeds, epochs 1–50 were compared by collapsed exact-entity validation AUPRC at fixed α = 0 and τ = 0.003. Scoring used unweighted L2-normalised residue vectors without an additional fixed-logit scale, and ties favored the earliest epoch. The selected checkpoints remained fixed during the validation parameter search described above. Final Fig. 5 evaluation used α = 0 and τ = 0.001 across all seeds, fractions and subsets. Each of the six single-vector runs was independently selected from epochs 1–50 by the same collapsed exact-entity validation AUPRC criterion, using its native pooled cosine score and resolving ties in favor of the earliest epoch. Validation AUPRC used all remaining cells in the collapsed validation candidate grid as operational negatives. The selected checkpoints and scoring parameters were held fixed for the database-filtered test evaluation described below. Final test encoding used per-sequence masks to exclude both terminal special tokens and padding before contextual encoding and pooling.

Protein–RNA test labels followed the same database-absence and taxonomy-prioritized sampling principle as PINDER. Identical inputs were collapsed, using the maximum score for duplicate entity pairs, to define a candidate grid of 459 proteins and 206 RNAs with 612 structural positives. Candidate negatives were excluded if their exact protein –RNA sequence pair occurred in the union of local BioLiP annotations and prepared-data audits, comprising 69,410 known edges.

Source-organism taxonomy was taken from PDB chain metadata, combining assignments across duplicate records. Structural positives were retained regardless of taxonomy; candidate negatives required taxonomy for both endpoints. Same-taxon pairs or ordered protein–RNA taxon combinations observed among positives were prioritized, followed by other taxonomically resolved pairs. Within each cohort, deterministic, score-independent ordering selected up to 100 negatives per positive. All methods shared the same labels within each cohort. The full, common source-excluded and sequence-distant sets contained 612, 576 and 254 positives with 61,200, 57,600 and 25,400 operational negatives, respectively. For the structure-distant set, all 12,881 eligible negatives were retained for 138 positives, giving a ratio of approximately 1 to 93.3. AUPRC and AUROC used these labelled pairs. Database absence denotes an operational negative rather than an experimentally verified non-interaction.

### Protein-RNA baselines

Graph-RPI used the released RPI1446 checkpoint, selected among five checkpoints by collapsed exact-entity validation AUPRC, whereas Transfer-RPI used the fixed released RPI2241 checkpoint; neither was retrained^11,33,39,40^. For the common comparison after exact source-sequence exclusion, exact raw-sequence matches to either checkpoint source were removed from every method, leaving a candidate grid of 447 proteins and 187 RNAs with 576 observed edges. Applying the shared negative-selection procedure yielded 58,176 labelled pairs, comprising 576 positives and 57,600 operational negatives. The external-method comparison, multi-vector versus single-vector comparison and PPI-data-fraction analysis all used this identical labelled set. RPISeq-RF and SeqMG-RPI were same-split retraining controls with one length-stratified corrupted negative per positive^41,42^. RPISeq-RF used RNA 4-mers and reduced-alphabet protein 3-mers; SeqMG-RPI used published RNA features, ESM-2 embeddings and predicted contact edges.^11,42^ Each same-split control was fitted and evaluated once, so no repeat-based uncertainty or significance comparison was assigned to these point estimates.

### AI-assisted preparation

AI-assisted tools supported language editing and analysis checks. Scientific interpretation and responsibility for the final manuscript, analyses and claims remain with the authors.

## Code availability

Code is available at https://github.com/UR-Free/ColBERT-PPI.

## Author contributions

H.Y. contributed to the experimental design, performed the computational experiments, analyzed the data and wrote the draft of the manuscript. R.L. analyzed the data and contributed to the interpretation and discussion of the results. J.Z. and Y.Z. designed the experiments, interpreted and discussed the results, and revised and improved the manuscript.

