## Supplementary Information for "Multi-vector retrieval enables residue-resolved prediction of protein partners by ColBERT-PPI"

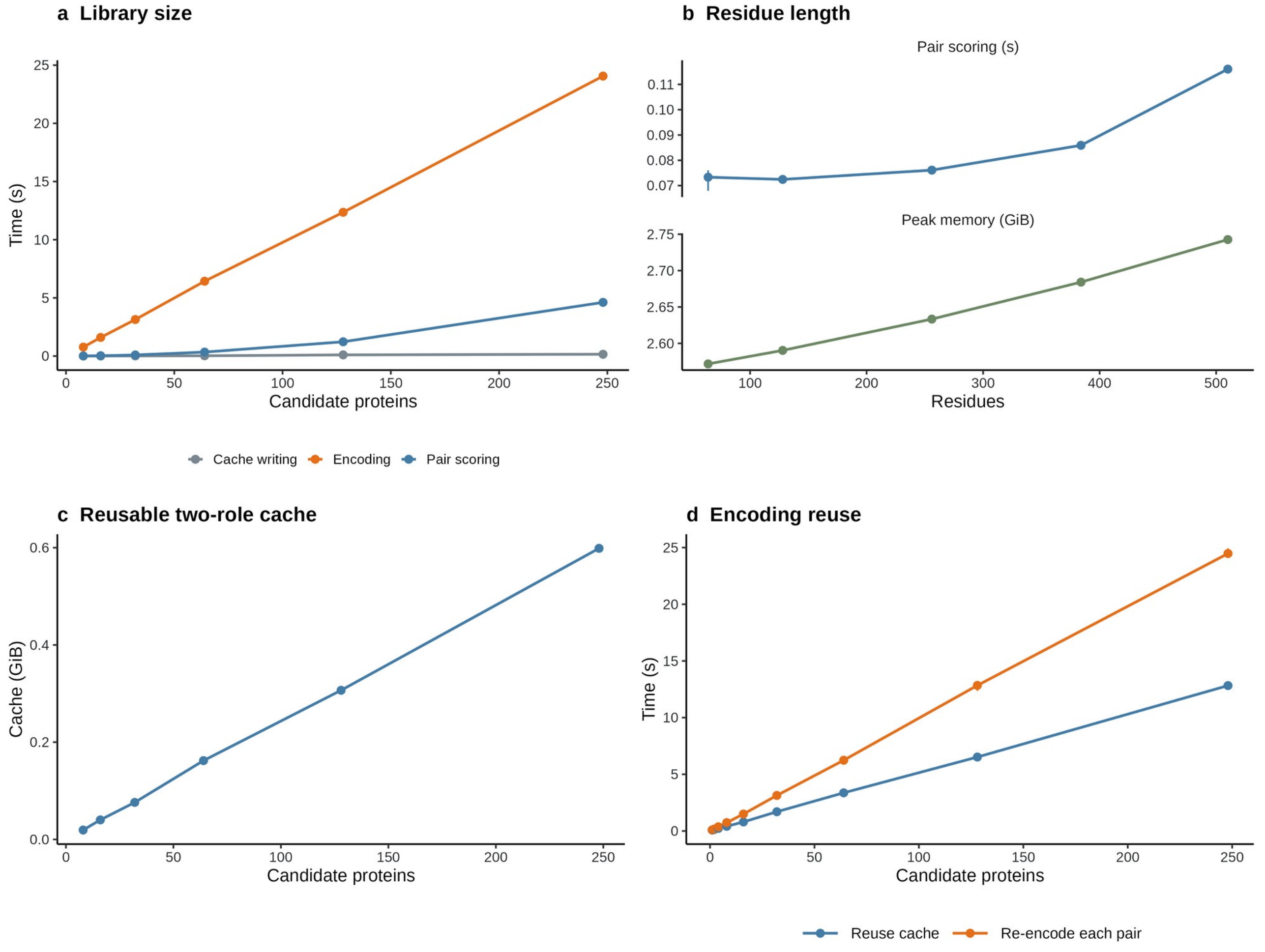


**Supplementary Fig. 1 | Empirical scaling and cache amortisation of ColBERT-PPI. a,** Encoding, cache materialisation and exact bidirectional scoring for N queries against N candidates, from 8 × 8 to 248 × 248 pairs. **b,** Scoring time and peak allocated memory for a fixed 32 × 32 matrix at residue lengths of 64 –510. **c,** Two-role FP32 cache size for the libraries in a, including per-residue background terms. **d,** Total time for one fixed query against 1–248 candidates, including encoding, cache materialisation and scoring, with monomers encoded once per role or re-encoded for every pair. The largest workloads in a and d therefore contain 61,504 and 248 pairs, respectively. All panels use calibrated smooth MaxSim ($\alpha = 1$, $\tau = 0.03$). Points are medians of three synchronised A100 measurements and ranges show minima and maxima.

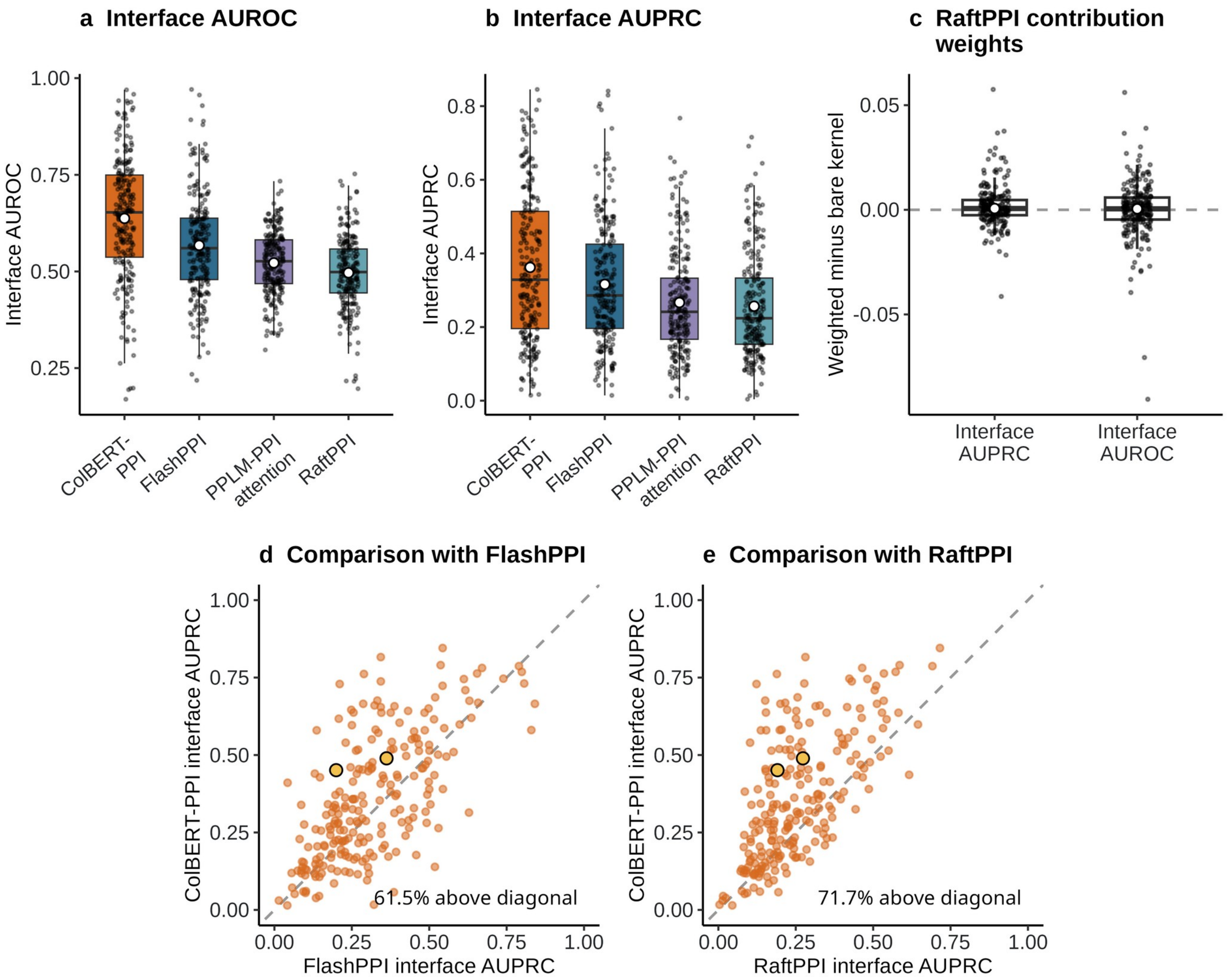


**Supplementary Fig. 2 | Regional interface evidence from each model's residue-pair scores. a-b,** Per-complex interface-residue AUROC (**a**) and AUPRC (**b**) across the fixed interface evaluation cohort (n = 226). ColBERT-PPI, FlashPPI and RaftPPI were trained on the PINDER training set; PPLM-PPI uses final-layer inter-protein attention from its released encoder. Boxes show medians and interquartile ranges, points show individual complexes and open circles show means. **c**, Paired difference between the weighted RaftPPI residue-contribution matrix and its unweighted Gaussian kernel; open circles show medians. **d-e**, Per-complex interface-residue AUPRC for ColBERT-PPI against FlashPPI (**d**) and RaftPPI (**e**). Diagonals denote equal performance, gold points mark the P62837 examples in **Figs. 3c, d** and labels give the fraction above the diagonal. No comparison required a fitted interpreter.

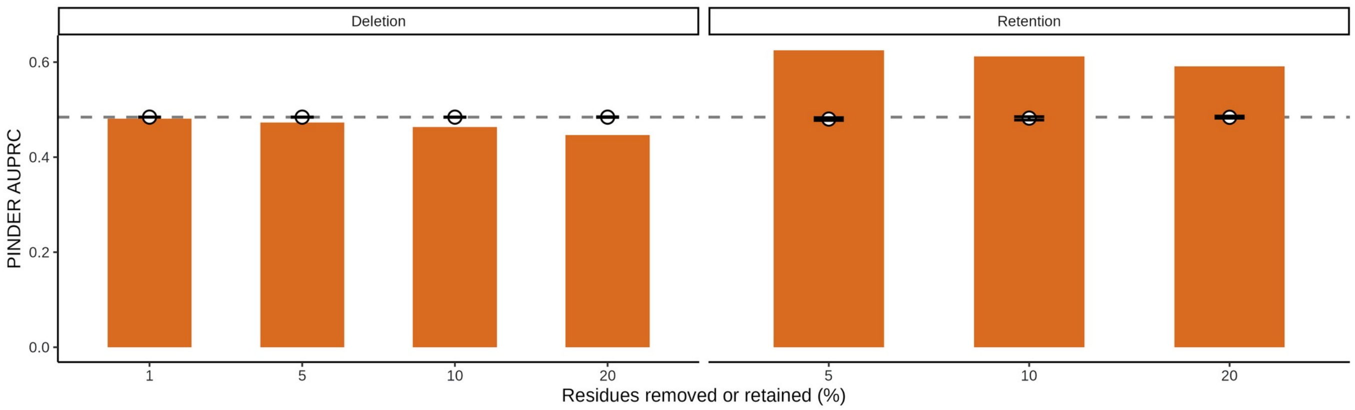


**Supplementary Fig. 3 | Score perturbations identify concentrated retrieval evidence.** PINDER AUPRC after removing the highest-contribution 1–20% of residues or retaining the highest-contribution 5–20% on the PINDER labelled test set. Bars show score-guided masks, open circles show means over 20 count-matched random masks, ranges show minima and maxima and dashed lines mark full-input AUPRC. Masks used frozen calibrated smooth MaxSim representations and background terms. Full-input AUPRC was 0.481726; the masks were defined using the observed partner and assess internal score concentration.

**Supplementary Table 1 | Training scale and evaluation status of PPI models.** Pair counts refer to different training stages and objectives. ColBERT-PPI, FlashPPI and RaftPPI used the PINDER training set; PPLM-PPI and PLM-interact used released checkpoints without retraining or PINDER checkpoint selection.

| Model | Pair-training source | Reported training scale | Training signal represented | PINDER evaluation status |
| --- | --- | --- | --- | --- |
| ColBERT-PPI | PINDER training set | 34,017 prepared records; 34,009 unique unordered accession pairs | Experimental residue contacts and sampled distant residue pairs | PINDER-trained; validation-selected epoch 69; calibrated smooth MaxSim |
| FlashPPI | PINDER training set | 34,017 prepared records | Protein-pair contrastive learning and residue-contact supervision | Trained on PINDER training set; validation-selected |
| RaftPPI | PINDER training set | 34,017 prepared records | Protein-pair retrieval | Trained on PINDER training set; validation-selected |
| PPLM-PPI | PDB and STRING paired pretraining; D-SCRIPT PPI heads | >3.3 million paired sequences in 672,372 clusters for paired pretraining | Paired masked-language modelling followed by supervised PPI heads | Released checkpoint; no retraining or PINDER selection |
| PLM-interact | Human STRING V11 | 421,792 pairs, comprising 38,344 positives and 383,448 negatives | Joint pair classification and masked-language modelling | Released checkpoint; no retraining or PINDER selection |

**Supplementary Table 2 | Complete-matrix inference measurements.** Each method was measured once after one warm-up on the PINDER timing matrix using A100-SXM4-80GB GPUs, FP32, TF32 off, a 510-residue cap and method-specific batches. Superscript a identifies PINDER-trained models and b identifies released-checkpoint evaluations. AUPRC was computed from the same 510-residue-cap score matrices over 24,543 labelled pairs (243 positives). ColBERT-PPI includes per-residue reference comparison in encoding time and the resulting background scalars in cache size; training-bank construction is offline. Total time uses an outer timer and can differ slightly from the sum of individually timed phases. Cache assembly denotes materialisation of encoded tensors for scoring.

| Method | AUPRC | Total time (s) | Encoding (s) | Cache assembly (s) | Pair scoring (s) | Peak allocated (GiB) | Reusable cache (GiB) |
|---|---|---|---|---|---|---|---|
| ColBERT-PPI[a] | 0.464906 | 29.724 | 24.629 | 0.223 | 4.867 | 3.84 | 0.599 |
| PPLM-PPI[b] | 0.098374 | 12018.682 | N/A | N/A | 12018.682 | 6.00 | N/A |
| FlashPPI[a] | 0.156234 | 22.521 | 22.516 | 0.002 | 0.002 | 3.28 | <0.01 |
| PLM-interact[b] | 0.064784 | 7626.574 | N/A | N/A | 7626.574 | 2.89 | N/A |
| RaftPPI[a] | 0.111582 | 1.031 | 1.021 | 0.005 | 0.004 | 0.84 | <0.01 |

**Supplementary Table 3 | Partner-conditioned residue masks preferentially suppress retrieval competitors.** Score reductions compare full input with the native-partner top-5% contribution-ranked residue mask across 243 PINDER systems. Intervals use 5,000 protein-endpoint bootstrap resamples. Negative reduction denotes a score increase after retention. Scores use calibrated smooth MaxSim ($\alpha = 1$, $\tau = 0.03$).

| Readout | Estimate | 95% interval |
|---|---|---|
| Native-partner score reduction | -0.0419 | -0.0468 to -0.0365 |
| Strongest-competitor score reduction | 0.0210 | 0.0148 to 0.0279 |
| Competitor minus native reduction | 0.0629 | 0.0553 to 0.0710 |
| Systems with greater competitor reduction | 98.8% | 95.6 to 100.0% |

**Supplementary Table 4 | Architecture ablations on sequence-distant and structure-distant PINDER and Y2H sets.** AUPRC and AUROC use all pairs in each fixed set. Bold indicates the highest point estimate and underlines the second highest within each dataset, set and metric. Complete and sequence-only multi-vector models use the same calibrated smooth MaxSim parameters with their respective training-reference banks; the single-vector model uses pooled cosine similarity.

| **Dataset** | **Method** | **Sequence-distant** | | **Structure-distant** | |
|---|---|---|---|---|---|
| | | **AUPRC** | **AUROC** | **AUPRC** | **AUROC** |
| **PINDER** | ColBERT-PPI | **0.659** | **0.969** | **0.609** | **0.966** |
| | Single vector | <u>0.545</u> | <u>0.946</u> | 0.494 | 0.939 |
| | Sequence only | 0.499 | 0.942 | <u>0.525</u> | <u>0.957</u> |
| **Y2H** | ColBERT-PPI | <u>0.385</u> | **0.726** | <u>0.424</u> | <u>0.734</u> |
| | Single vector | 0.288 | 0.664 | 0.310 | 0.664 |
| | Sequence only | **0.479** | <u>0.716</u> | **0.527** | **0.747** |

**Supplementary Table 5 | External-method comparison on sequence-distant and structure-distant Y2H sets.** AUPRC and AUROC use all labelled pairs in each fixed set. Bold indicates the highest point estimate and underlines the second highest within each set and metric.

| **Method** | **Sequence-distant** | | **Structure-distant** | |
|---|---|---|---|---|
| | **AUPRC** | **AUROC** | **AUPRC** | **AUROC** |
| ColBERT-PPI | **0.385** | **0.726** | **0.424** | **0.734** |
| FlashPPI | 0.259 | 0.577 | 0.301 | 0.596 |
| PPLM-PPI | <u>0.268</u> | <u>0.637</u> | <u>0.317</u> | <u>0.640</u> |
| PLM-interact | 0.213 | 0.557 | 0.242 | 0.562 |
| RaftPPI | 0.200 | 0.463 | 0.217 | 0.446 |
